# Expression pattern of the ST heat-stable enterotoxin gene cassettes in *Vibrios* superintegrons

**DOI:** 10.1101/2024.10.24.620042

**Authors:** Érica L. Fonseca, Fernanda S. Freitas, Sérgio M. Morgado, Ana Carolina P. Vicente

**Affiliations:** Laboratório de Genética Molecular de Microrganismos, Instituto Oswaldo Cruz, FIOCRUZ, Rio de Janeiro, Brazil

**Author notes:** Corresponding Author. Érica L. Fonseca. Avenida Brasil 4365, Manguinhos, Rio de Janeiro, 21040-360, Brazil.

**Keywords:** virulence, gene expression regulation, chromosomal integron, cholera-like diarrhea, relative quantification, transcription level

## Abstract

**BACKGROUND:** *Vibrio mimicus* has been associated with a cholera-like diarrhea and other gastroenteritis due to the presence of virulence factors such as the heat-stable ST enterotoxin. In spite of its relevant role in *Vibrio* pathogenesis, there is still a lack of information concerning ST expression dynamics and associated genomic context.

**OBJECTIVE:** this study aimed to determine the prevalence, genomic context, and expression of the *st* gene found in *V. cholerae* and *V. mimicus* strains.

**METHODS:** genome sequencing, in silico analyses of st promoter region and genomic context, and relative quantification pf st expression by real time RT-PCR were applied.

**FINDINGS:** The *st* gene was found within the SI of both *Vibrio* species, presenting variations in their amino acid sequences. The *in silico* analysis of the *st* 5’UTR identified different promoter versions among them. Significant differences were observed in the *st* expression level, which varied from 10X to 640X-fold, correlating with the differences found in the promoter versions.

**MAIN CONCLUSIONS:** Therefore, the present study highlighted the link between the ST toxin gene and the superintegron, not only for *V. cholerae*, but also for *V. mimicus*. The differential *st* expression revealed that it is detrimental to the cognate promoter version, which would impact the toxin production and strain virulence.

---

*Vibrio mimicus* is a ubiquitous organism in fresh and sea waters and has been associated with cholera-like diarrhea and other gastroenteritis.^(1)^ Some *V. mimicus* strains may harbor the *ctx* and *tcp* operons, which encode the cholera enterotoxin (CT) and the toxin co-regulated pilus (TCP), the main virulence factors involved in choleragenic *V. cholera*.^(2)^ However, *V. mimicus* strains lacking these virulence determinants have also been recovered from diarrhea outbreaks.^(3)^ The heat-stable enterotoxin (ST) coded by the *st* gene is another virulence factor occurring in Vibrios causing diarrhea.^(4-6)^ It encodes a 17-amino acid peptide and shares a high identity with the ST produced by enterotoxigenic *Escherichia coli* (ETEC).^(7)^ Two major *st* alleles have been characterized in *V. cholerae* NAG and *V. cholerae* O1 strains (named *stn* and *sto* that code the ST-NAG and ST-O1, respectively).^(6)^ Further, it was verified that the *stn* gene was prevalent in *V. mimicus* strains from both clinical and environmental origin, and some authors suggested that this species would be the reservoir of this gene. ^(4,8)^ Interestingly, independent of the *Vibrio* species, it was demonstrated that both *stn* and *sto* genes were flanked by 123-bp almost identical direct repeats,^(6)^ which were subsequently characterized to be the recombination sites in the *V. cholerae* superintegron (SI).^(5,6,9,10)^ The SIs are chromosomal genetic platforms carrying up to hundreds of gene cassettes, which can easily be inserted, excised, or rearranged within the SIs.(11,12)

Despite the clinical relevance and the prevalence of ST toxin, its functionality had only been indirectly demonstrated through the suckling model and ELISA assays. ^(6,13)^ Moreover, until now, the dynamics of *stn* and *sto* expression in both *V. cholerae* and *V. mimicus* have never been assessed. To contribute to this issue, we characterized the genomic context of the *stn* gene and evaluated its functionality by assessing its expression in environmental and clinical *V. mimicus* and *V. cholerae* strains.

## MATERIAL AND METHODS

*Bacterial strains -* During a genomic study on Vibrios from our Bacterial Culture Collection, it was verified the presence of the *stn* gene in genetically unrelated *V. cholerae* NAG and *V. mimicus* strains.^(14)^ Three *V. cholerae* NAG strains (VC688, VC712 and VC716) were recovered from human and animal in Japan, while the environmental *V. mimicus* strains (VM602, VM605, and VM606) were isolated from water in Brazil.^(5,14)^ The *stn* genes in each species were named from now on, *stn-vc* for the *V. cholerae* NAG and *stn-vm* for the *V. mimicus* strains.

*Genome Sequencing -* The genomic DNA extraction was performed using the NucleoSpin Microbial DNA kit (Mach-erey Nagel), and the genomic libraries were constructed using Nextera paired-end library. The sequencing was performed using Illumina Hiseq 2500, generating reads of 150 bp length. The raw reads were filtered and trimmed using NGS QC Toolkit v.2.3.3 with a Phred score ≥ 20. The genomes were *de novo* assembled using SPAdes assembler v3.14.1.^(15)^ The genomic context of *stn* genes was evaluated and compared with each other as well as their deduced amino acid sequences. An *in silico* promoter prediction analysis was performed to identify the putative promoters of these *stn* genes, and the predicted promoters were compared with those previously identified for *stn* and *sto* genes.^(6,16)^ The genomes of the ST-positive *V. cholerae* and *V. mimicus* strains analyzed here were deposited in GenBank under accession nos. JBDFMU000000000 (VC688), JBDFMT000000000 (VC712), JBDFMS000000000 (VC716), JBAKBX000000000 (VM602), JBAKBW000000000 (VM605), JAZHPS000000000 (VM606).

*Relative Quantification of ST toxin expression -* To assess whether the *stn-vc* and *stn-vm* identified here are expressed, the total RNA was isolated using the NucleoSpin RNA Isolation kit, according to the manufacturer”s instructions. Before cDNA synthesis, RNA extracts were treated with Turbo DNA-free reagent (Ambion, Applied Biosystems), according to the supplier”s recommendations, to eliminate contaminant genomic DNA. After DNase treatment, RNA samples were quantified in a Qubit RNA HS Assay Kit (thermoFisher) and diluted to the same concentration (10 ng/mL) with sterile nuclease-free water. Total RNA extracts were submitted to cDNA synthesis with SuperScript III RNase H-reverse transcriptase (ThermoFisher) and the resulting cDNA was used as a template in real-time PCR assays.

The transcription of the *stn* genes was evaluated by real-time RT-PCR using the Power-SYBR Green PCR Master Mix (ThermoFisher). The ST expression of both *V. mimicus* and *V. cholerae* was assessed with the primers ST RT F 5” AACAGTGCAGCAACCACAAC 3”, and ST RT R 5” TGGGCATTCTTCATTTTCAC 3”. In order to correct for differences in the amount of starting RNA, the single-copy housekeeping *rpoA* genes of *V. cholerae* (VC RPOA RT F 5” GAACAAATCAGCACGACACA 3”, and VC RPOA RT R 5” CACAACCTGGCATTGAAGA 3”) and *V. mimicus* (VM RPOA RT F 5” GTGGTCGTGGTTATGTTCCAG 3”, and VM RPOA RT R 5” CGCTCGTCTTCTTCAGTATGG 3”) were used as reference genes for normalizing the RQ.

Independent assays were performed in triplicate to determine the reliability of the relative quantification (RQ). The RQ results were presented as ratios of gene transcription between the target gene (*stn-vm* and *stn-vc*) and the reference gene (*rpoA-vm* and *rpoA-vc*), which were obtained according to the following equation: RQ=2^-ΔCT^, where CT is the value corresponding to the crossing point of the amplification curve with the threshold and ΔCT=CT target gene - CT reference gene.

To evaluate the statistical significance of the differences in the expression levels observed among the strains, an analysis of variance (ANOVA) was applied followed by the Tukey test for a single-step multiple comparison of means, in which a *p* < 0,05 value was considered statistically significant.

## RESULTS AND DISCUSSION

*Analysis of ST-VM and ST-VC amino acid sequences -* Analysis of the predicted amino acid sequence of *stn-vc* (n=3) and *stn-vm* (n=3) from this study by BLASTp revealed some slight differences among them. We identified five differences among the ST-VM and ST-VC in the first 27 residues. The ST of the VC688 and VC716 NAG strains were identical to the ST-NAG previously reported.^(16)^ The exception was VC712, which had a substitution at residue 27 that was not present in the canonical ST-NAG sequence. On the other hand, the ST-VM sequences shared amino acid residues with both ST-O1 and ST-NAG (Table 1). The ST is encoded as a preprotoxin, comprising a signal sequence, a mature sequence, and a central region between them. The signal sequence consists of the first 18 amino acids with a hydrophobic character and is involved with the protection of the mature toxin from endogenous proteolytic action until its cleavage upon export to periplasm.^(17)^ Therefore, the modifications observed here occurred in the signal and central sequences (Table 1) with no impact on the mature ST toxin, as observed previously.^(6)^

**Table 1.**
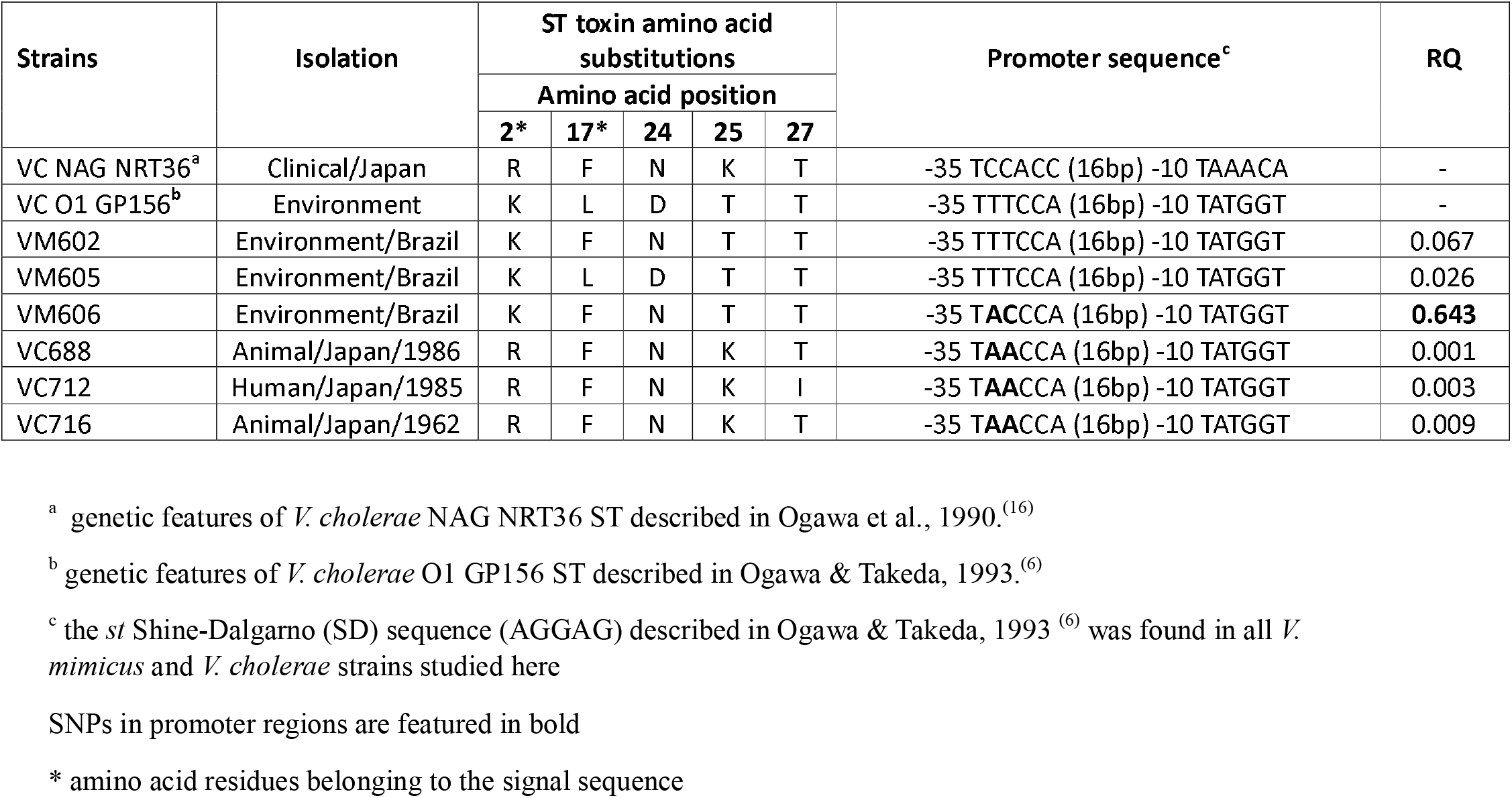
Differences in the amino acid and promoter sequences of the ST-VM and ST-VC and their corresponding expression levels of the *V. cholerae* and *V. mimicus* strains analyzed in this study

*Genomic context of stn-vm and stn-vc and characterization of promoter regions -* The genomic analysis revealed that the *stn-vc and stn-vm* genes presented in clinical *V. cholerae* NAG and environmental *V. mimicus* strains, respectively, were a single gene within the SI, and flanked by the 123-bp repeats.^(5,6)^ Considering that the association of *stn* with direct repeats/recombination sites in the context of SIs may favor its dispersion in the genus *Vibrio*,^(5,6,9,10)^ we searched for the *stn*/*sto* genes in the 10,117 *V. cholerae* and 44 *V. mimicus* genomes available in GenBank. We found 8 ST-positive *V. cholerae* (most NAGs) and four ST-positive *V. mimicus*, all of them in the context of SIs in different lineages (data not shown), showing the potential mobility of the *st* gene.

The finding regarding the occurrence of both *stn-vc* and *stn-vm* in SIs raised the question of whether these genes were expressed. Since genes encoded in distal positions within the SIs are usually not expressed due to their physical distance from the SI promoter region, the efficient expression of *stn* genes would be dependent on their relative position in the SIs.^(11,18)^ The ST production had already been indirectly demonstrated through the suckling model and ELISA,^(6,13)^ indicating that the *st* gene is expressed in those strains. In fact, it had been demonstrated the presence of promoter regions and Shine-Dalgarno sequences in the 5”UTR of *sto* and *stn* genes of *V. cholerae* O1 and NAG strains, respectively.^(6,16)^ The ST-O1 (*sto*) promotor region (−35 TTTCCA [16 bp] -10 TATGGT) was different from the one described for ST-NAG (*stn*) (−35 TCCACC [16 bp] -10 TAAACA). Therefore, we searched for both promotor regions in the 5”UTR of the *stn*-*vc* and *stn*-*vm* genes from this study. This analysis revealed the presence of putative promoters for *stn* occurring in *V. mimicus*, which had never been demonstrated. Moreover, a diversity of promoter sequences among *stn-vc* and *st-vm* genes was verified. The *st* promoter region of the *V. cholerae* NAG strains from our study was identical to each other but different from both ST-O1 and ST-NAG promoters previously described (Table 1).^(6,16)^ Interestingly, although VC688, VC712 and VC716 were all *V. cholerae* NAG strains, their promoter resembled the one found in *V. cholerae* O1, since they shared the same -10 TATA box and presented two polymorphisms in -35 hexamer (−35 T**AA**CCA [16bp] -10 TATGGT).^(6)^ On the other hand, while VM602 and VM605 presented *stn-vm* promoter sequences identical to that of ST-O1, the VM606 presented another promoter configuration with two polymorphisms in -35 box (−35 T**AC**CCA [16bp] -10 TATGGT) (Table 1). Therefore, these results indicate that the *stn-vc* and *stn-vm* genes found here carry a promoter and may be efficiently expressed independently of the SI promoter.

*Relative expression of stn-vm and st-vc -* To verify the impact of these different putative promoter versions on ST expression, their functionality was assessed by a relative quantification (RQ) assay. The RQ analysis revealed that the *stn* was expressed in all strains independently of the promoter version.

The three *V. cholerae* NAG strains, besides harboring the same promoter configuration, presented similar transcription levels (RQ values VC688=0.001, VC712=0.003, VC716=0.009). On the other hand, the *V. mimicus* strains presented discrepancies in the *stn-vm* transcription (RQ values VM602=0.067, VM605=0.026, VM606=0.643).

Interestingly, we observed a correlation of the *stn-vc* and *stn-vm* expression level with the promoter version identified. The *stn-vm* of VM606, which harbored a unique promoter configuration was, by far, the most expressed, with a 10X-to 25X-fold increase in expression relative to the *st*-*vm* of VM602 and VM605, respectively, both harboring the canonical ST-O1 promoter (Table 1). Considering the *V. cholerae* strains, these differences were even more remarkable, in which *st-vm* of VM606 was 70X-to 640X-fold more expressed than *st*-*vc* genes, which presented another promoter version (Table 1). Taking into account that the *stn-vc* and *stn-vm* genes were in a single copy in *V. cholerae* and *V. mimicus* genomes analyzed here, these findings strongly indicated that the polymorphisms found in the promoter regions of *stn-vm* and *stn-vc* (Table 1) were directly involved with the differential expression observed. In this way, the unique promoter version found in *stn-vm* of VM606 would confer the most efficient transcription apparatus considering the other ones identified, suggesting that the polymorphisms in the -35 hexamer may directly affect the transcription efficiency.

The alterations in the gene expression patterns due to point mutations in the promoter region have already been demonstrated. A study verified that the expression of *bla*_NDM-5_ suffered a significant decrease due to one point mutation in its ribosomal binding site,^(19)^ while in another study, a unique point mutation in the -35 hexamer of the class 1 integron Pc promoter was responsible for the overexpression of *bla*_IMP-5_ gene.^(20)^

To check for the distribution of the ST promoter versions identified here, we searched for the promoter region in the *stn/sto* genes with available 5”UTR found in *V. cholerae* (n=5) and *V. mimicus* (n=4) sequences in GenBank. Interestingly, it was verified that any of them harbored the ST promoter version found in VM606 strain, the one with the highest ST expression values (Table 1). Instead, 4/9 *st* sequences (n= 2 *V. cholerae* and n=2 *V. mimicus*) harbored the promoter version found in the three *V. cholerae* NAG strains from this study, which presented the lowest ST expression levels. Two of them (2/9; n=1 *V. cholerae* and n=1 *V. mimicus*) were identical to the promoter previously described for the ST-O1^(6)^ that was also found in the *stn-vm* of VM602 and VM605, with intermediate expression levels (Table 1). Moreover, three new promoter versions were identified, two of them presented polymorphisms in the -35 hexamer (−35 T**GA**ACA and -35 T**AG**ACA), both occurring in the *st* gene from *V. cholerae*, including one (−35 T**AG**ACA) that was recovered from a human case in USA/2023. The other version was found in the *st* of one *V. mimicus* (−35 **CTT**CCA [16 bp] -10 T**G**TGGT).

This heterogeneity of promoters suggests that both *stn* and *sto* genes occurring in *V. mimicus* and *V. cholerae* may present different spectra of expression, which would reflect different levels of toxin production and strain virulence.

## CONCLUSION

Therefore, the SIs may contribute to the virulome of *V. cholerae* and *V. mimicus* and, at least for the *st* gene, the expression of this virulence factor is dependent on its cognate promoter version and assured irrespective of its genomic context.

## ACKNOWLEDGMENTS

To Nathália M. S. Bighi for the support with statical analysis.

## FUNDING

This work was supported by Conselho Nacional de Desenvolvimento Científico e Tecnológico (CNPq) and Oswaldo Cruz Institute grants.

## AUTHOR’S CONTRIBUTION

**ELF:** Data curation, Visualization, Validation, Methodology, Writing-Original draft preparation, Writing - Review & Editing. **SMM:** Methodology, Visualization, Formal analysis, Data curation, Writing - Review & Editing. **FSF:** Methodology. **ACPV:** Conceptualization, Writing - Review & Editing, Validation, Funding acquisition, Supervision.

## CONFLICT OF INTEREST

None to declare

## Notes

### Competing Interest Statement

The authors have declared no competing interest.

## REFERENCES

1. Baker-Austin C, Oliver JD, Alam M, Ali A, Waldor MK, Qadri F, Martinez-Urtaza J. Vibrio spp. Infections. Nat Rev Dis Primers. 2018; 4:8. doi:10.1038/s41572-018-0005-8.

2. Faruque SM, Albert MJ, Mekalanos JJ. Epidemiology, genetics, and ecology of toxigenic Vibrio cholerae. Microbiol Mol Biol Rev. 1998; 62:1301–1314. doi:10.1128/MMBR.62.4.1301-1314.1998.

3. Alam MT, Stern SR, Frison D, Taylor K, Tagliamonte MS, Nazmus SS, et al. Seafood-Associated Outbreak of ctx-Negative Vibrio mimicus Causing Cholera-Like Illness, Florida, USA. Emerg Infect Dis. 2023; 29:2141–2144. doi: 10.3201/eid2910.230486.

4. Yuan P, Ogawa A, Ramamurthy T, Nair GB, Shimada T, Shinoda S, et al. Vibrio mimicus are the reservoirs of the heat-stable enterotoxin gene (nag-st) among species of the genus Vibrio. World J Microbiol Biotechnol. 1994;10:59–63. doi: 10.1007/BF00357565.

5. Vicente AC, Coelho AM, Salles CA. Detection of Vibrio cholerae and V. mimicus heat-stable toxin gene sequence by PCR. J Med Microbiol. 1997;46:398–402. doi: 10.1099/00222615-46-5-398.

6. Ogawa A, Takeda T. The gene encoding the heat-stable enterotoxin of Vibrio cholerae is flanked by 123-base pair direct repeats. Microbiol Immunol. 1993;37:607–616. doi: 10.1111/j.1348-0421.1993.tb01683.x.

7. Arita M, Takeda T, Honda Y, Miwatani T. Purification and characterization of Vibrio cholerae non-O1 heat-stable enterotoxin, Infect Immun. 1986;52:45–49. doi: 10.1128/iai.52.1.45-49.1986.

8. Yuan P, Ogawa A, Ramamurthy T, Nair GB, Shimada T, Shinoda S, et al. Vibrio mimicus are the reservoirs of the heat-stable enterotoxin gene (nag-st) among species of the genus Vibrio. World J Microbiol Biotechnol. 1994;10:59–63. doi: 10.1007/BF00357565.

9. Takeda T, Peina Y, Ogawa A, Dohi S, Abe H, Nair GB, et al. Detection of heat-stable enterotoxin in a cholera toxin gene-positive strain of Vibrio cholerae O1. FEMS Microbiol Lett. 1991;64:23–27. doi: 10.1016/0378-1097(91)90203-m.

10. Rowe-Magnus DA, Mazel D. Integrons: natural tools for bacterial genome evolution. Curr Opin Microbiol. 2001;4:565–569. doi: 10.1016/s1369-5274(00)00252-6.

11. Mazel D, Dychinco B, Webb VA, Davies J. A distinctive class of integron in the Vibrio cholerae genome. Science. 1998;280:605–608. doi: 10.1126/science.280.5363.605.

12. Hasan NA, Grim CJ, Haley BJ, Chun J, Alam M, Taviani E, et al. Comparative genomics of clinical and environmental Vibrio mimicus. Proc Natl Acad Sci U S A. 2010;107:21134–21139. doi: 10.1073/pnas.1013825107.

13. Nair GB, Bhattacharya SK, Takeda T. Identification of heat-stable enterotoxin-producing strains of Yersinia enterocolitica and Vibrio cholerae non-O1 by a monoclonal antibody-based enzyme-linked immunosorbent assay. Microbiol Immunol. 1993;37:181–186. doi: 10.1111/j.1348-0421.1993.tb03198.x.

14. Vieira VV, Teixeira LF, Vicente AC, Momen H, Salles CA. Differentiation of environmental and clinical isolates of Vibrio mimicus from Vibrio cholerae by multilocus enzyme electrophoresis. Appl Environ Microbiol. 2001;67:2360–2364. doi: 10.1128/AEM.67.5.2360-2364.2001.

15. Bankevich A, Nurk S, Antipov D, Gurevich AA, Dvorkin M, Kulikov AS, et al. SPAdes: a new genome assembly algorithm and its applications to single-cell sequencing. J Comput Biol. 2012;19:455–77. doi: 10.1089/cmb.2012.0021.

16. Ogawa A, Kato J, Watanabe H, Nair BG, Takeda T. Cloning and nucleotide sequence of a heat-stable enterotoxin gene from Vibrio cholerae non-O1 isolated from a patient with traveler”s diarrhea. Infect Immun. 1990;58:3325–3329. doi: 10.1128/iai.58.10.3325-3329.1990.

17. Nair GB, Takeda Y. The heat-stable enterotoxins. Microb Pathogen. 1998;24:123–131. Doi: 10.1006/mpat.1997.0177.

18. Cambray G, Sanchez-Alberola N, Campoy S, Guerin E, Da Re S, González-Zorn B, et al. Prevalence of SOS-mediated control of integron integrase expression as an adaptive trait of chromosomal and mobile integrons. Mob DNA. 2011;2:6. doi: 10.1186/1759-8753-2-6.

19. Li Y, Zhang R, Wang S. A Natural Novel Mutation in the bla(NDM-5) Promoter Reducing Carbapenems Resistance in a Clinical Escherichia coli Strain. Microbiol Spectr. 2022;10:e0118321. doi: 10.1128/spectrum.01183-21.

21. Brízio A, Conceição T, Pimentel M, Da Silva G, Duarte A. High-level expression of IMP-5 carbapenemase owing to point mutation in the-35 promoter region of class 1 integron among Pseudomonas aeruginosa clinical isolates. Int J Antimicrob Agents. 2006;27:27–31. doi: 10.1016/j.ijantimicag.2005.08.023.

